# HIP IMO Report: Analyzing Phenotypic Properties of Bladder Cancer Using Ordinary Differential Equation (ODE) Models

**DOI:** 10.1101/839209

**Authors:** Benjamin Sherwin

**Affiliations:** H. Lee Moffitt Cancer Center and Research Institute

## Abstract

Bladder cancer is composed of proliferative and immunogenic phenotypes, which ultimately play a significant role in the growth of the tumor. By using ordinary differential equation models, this paper models the impact of high and low immunogenic cell populations on non-muscle invasive bladder cancer when treated with and without the Bacillus Calmette-Guerin vaccine. Furthermore, this paper models the impact that the Bacillus Calmette-Guerin vaccine has on inflammatory cytokines, which inhibit the growth of tumors by stimulating an immune response. We focus primarily on how the immunogenicity phenotype impacts population dynamics in non-muscle invasive bladder cancer.

## I. Introduction to bladder cancer

Bladder cancer is the ninth most diagnosed malignancy worldwide, yet it is relatively understudied in oncology research. Therefore, it is crucial that the scientific community utilize resources like quantitative and experimental data to help patients increase their probabilities of survival.

Furthermore, bladder cancer is divided into two different types based on its severity: non-muscle invasive and muscle invasive. Non-muscle invasive bladder cancer (NMIBC) can progress into muscle invasive bladder cancer (MIBC), which is more deadly with a five-year mortality rate of approximately 50-70 percent. MIBC has higher recurrence rates and poor prognosis, which leads to further problems for patients. This includes a procedure to remove the entire bladder, called a radical cystectomy, which is used to treat MIBC. For NMIBC, the standard treatment is a vaccine called Bacillus Calmette Guerin (BCG). Currently, BCG treatment is the best available option to stop the fatal progression from NMIBC to MIBC.

BCG is an intravesical immunotherapy that is typically utilized to fight against tuberculosis in children but can also be utilized to treat NMIBC by stimulating an immune response. This is because the vaccine contains the *Mycobacterium tuberculosis* bacteria which is subsequently released into the tumor. Thus, the lymphocytes in the immune system produce antibodies, which fight against the antigen. Inadvertently, this activated immune response also affects the cancer cells in response to the BCG vaccine. The BCG course of treatment consists of three key phases: induction, maintenance, and follow-up. In the induction phase, patients receive weekly instillations of the vaccine until there is a reduction in tumor size, which usually takes about six weeks. Furthermore, the maintenance phase consists of weekly instillations for about three weeks at specified intervals. In addition, urine cytology is tested at an interval of three weeks. After about 24 weeks, the patient will enter the follow-up phase which involves urine cytology tests every six weeks. Unfortunately, BCG treatment only has a success rate of about 58% with 13% of patients progressing from NMIBC to MIBC.

## II. Bladder cancer phenotypes at the cellular level

Bladder cancer tumors are made up of a variety of different cells, each displaying a different phenotype. The two most common phenotypes displayed in these tumors are immunogenic and proliferative cells. The composition of a bladder cancer tumor is heterogenous in nature, consisting of varying degrees of proliferative and immunogenic cells, as shown in Figure 1A. For the purpose of this project, the focus was on high and low immunogenic cells rather than high and low proliferative cells.

**Figure 1.**
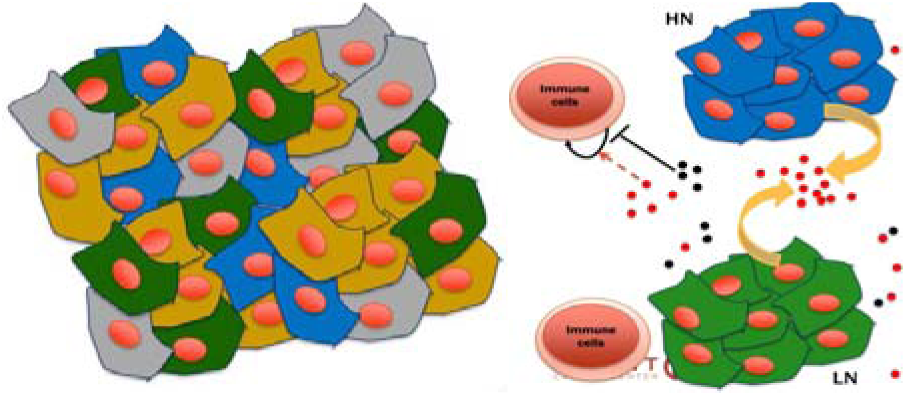
A: Bladder cancer tumor composed of four different phenotypic cell types: low proliferative, high proliferative, low immunogenic, and high immunogenic B: Diagram of the interactions between immune and tumor cells, where HN is high immunogenic and LN is low immunogenic. The red and black dots represent suppressants and anti-suppressants released by immune cells that interfere with tumor cells.

As shown in Figure 1B, it appears that immune response does play a fundamental role in bladder cancer tumor growth by releasing both anti-inflammatory and inflammatory cytokines. Cytokines are proteins secreted by immune cells that serve as chemical messengers for the immune system. This causes immune cells to trigger mechanisms to respond to invaders in the body. BCG treatment causes an immune response which results in the release of these cytokines at the site of the tumor, making it an effective immunotherapy. The inflammatory cytokines that are released by the immune cells causes increased death of bladder cancer cells. It is important to note that cytokines cause high immunogenic cells to die more rapidly than low immunogenic cells.

According to Helegaard et al, immunogenicity phenotypes affect the survival probability of a patient with a p-value of 0.0016, proving its statistical significance. As shown in Figure 2A, high immune response is associated with consistently higher survival probabilities than low immune response over time. This indicates that immune response is associated with better outcomes in MIBC. Moreover, Figure 2B illustrates the connection between mutation levels and immune response. As mutation burden increases, immune response also increases in a direct relationship. This can be seen by the increase in median values on the box plots in the figure. Thus, it is assumed that by increasing mutation burden and immune response, survival probability will also increase.

**Figure 2.**
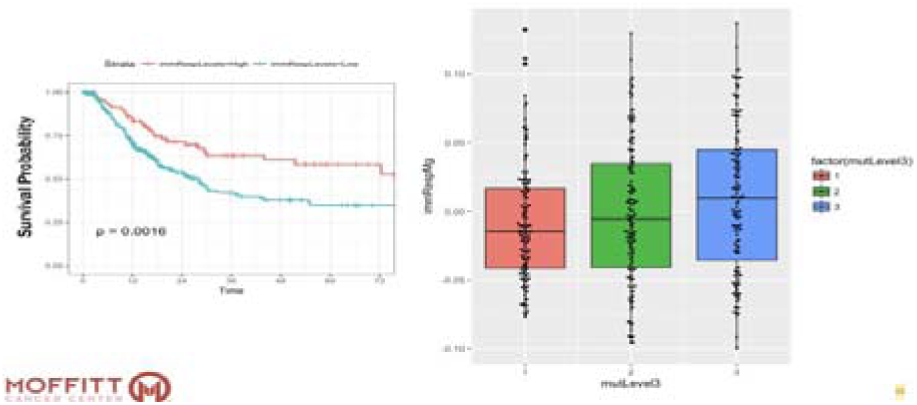
A: The graph reveals that a high immune response results in a higher survival probability for the patient than that of a low immune response B: The box plot illustrates that when the level of mutations increase, immune response also increases leading to higher survival probabilities.

In addition, mutation burden can refer to an increase in immunogenic mutations causing the tumor to display a highly immunogenic phenotype. As shown Figure 3 by Helegaard et al, increased mutation burden correlates with higher survival probabilities specifically in highly proliferative tumors. High mutation rates are associated with higher survival probabilities specifically in highly proliferative tumors. High mutation rates are associated with high survival probability over time as indicated by the graph. The p-value for this relationship is 0.00011, revealing its statistical significance. Thus, there may be a correlation between immunogenicity and survival probability, which could potentially help patients survive bladder cancer.

**Figure 3.**
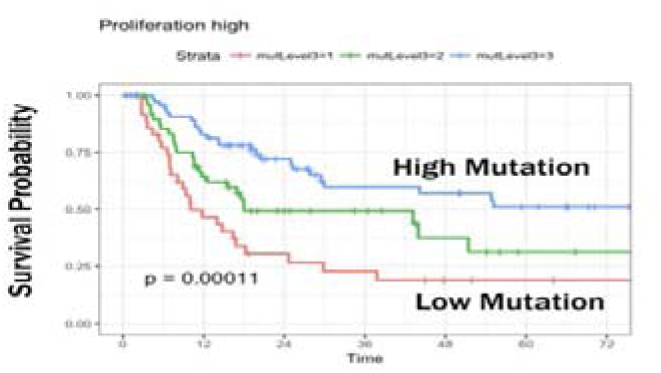
The graph reveals that a high mutation burden results in a higher survival probability for the patient than that of a low mutation burden with a p-value of 0.00011

## III. Bladder cancer ordinary differential equation model

### A. Hypotheses

Key hypotheses that are essential to the construction of the ODE model include:

1. Phenotypes play a role in treatment and tumor growth
2. If there is an increased amount of inflammatory cytokines from the BCG treatment, then tumor populations decrease.

### B. Equations and Parameters

The following mathematical equations model the population dynamics of low immunogenic cells, high immunogenic cells, and inflammatory cytokines. Parameters for each of these ordinary differential equations are crucial to understanding the growth of bladder cancer tumors both in the presence and not in the presence of the BCG treatment.

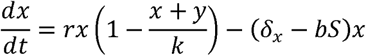

The differential equation above models the change in population of low immunogenic cells over time. The first part of the equation, *rx*, characterizes the growth rate of the tumor where r is the rate of growth and x is the initial number of low immunogenic cells. The following term, 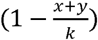, is used to prevent unregulated exponential growth by setting a carrying capacity, *K*. Finally, the subtraction of the last term represents the death of the tumor cells where *δ*_*x*_ represents the death rate of low immunogenic cells, b represents death cause by cytokines released by the immune system, and S represents the concentration of cytokine. This is multiplied by the number of low immunogenic cells to account for the entirety of the tumor.

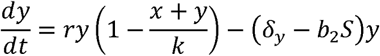

The differential equation above represents the change in population of high immunogenic cells over time. This equation is very similar to that of the low immunogenic cells with all the terms serving the same purpose. However, there are some significant differences. When incorporating the first term for growth and the last term for death, y is used instead of x since it represents high immunogenic cells. Furthermore, *δ*_*y*_ and *b*_*2*_ have subscripts so that the parameters can be changed to account for high immunogenic cells. This is important to include because the death rate and death induced by cytokines vary based on immunogenicity phenotypes.

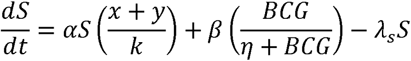

The differential equation above represents the change in the concentration of inflammatory cytokines over time. The first term 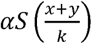 sets a carrying capacity for logistic growth and is regulated by the concentration of cytokines, S, as well as a scaling term, *α*. The second term, 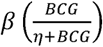, accounts for the BCG treatment for the increased release of cytokines by the immune system, with the scaling term *β*. Furthermore, the term 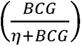 is a Hill equation used to control the effects of BCG on the tumor. Finally, the last term accounts for the decline in inflammatory cytokine concentration with a decay rate *λ*_*S*_, multiplied by the overall concentration of cytokines in the tumor.

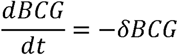

The differential equation above represents the decay of the BCG vaccine over time, where -*δ* is the decay rate and BCG is the amount of vaccine given to a patient. The equation helps to replicate what occurs in the tumor since the BCG has a weaker effect as time increases from when it was administered.

### C. Model Exploration and Graphs

*Note: All graphs showing tumor populations use blue for low immunogenicity, red for high immunogenicity, and yellow for total tumor growth. Graphs with only a single line of blue represent cytokine concentration. In addition, all models were produced using MATLAB*.

In Figures 4A and 4B below, the graphs model both the inflammatory cytokines and tumor phenotype populations without BCG treatment. In Figure 4B, the model reveals the normal growth of a bladder cancer tumor without any treatment over a period of about five years. As shown, the low immunogenic cells are able to outcompete the high immunogenic cells causing the high immunogenic population to decrease more rapidly. In addition, the tumor follows a logistic growth curve in this case but may vary in real life.

**Figure 4.**
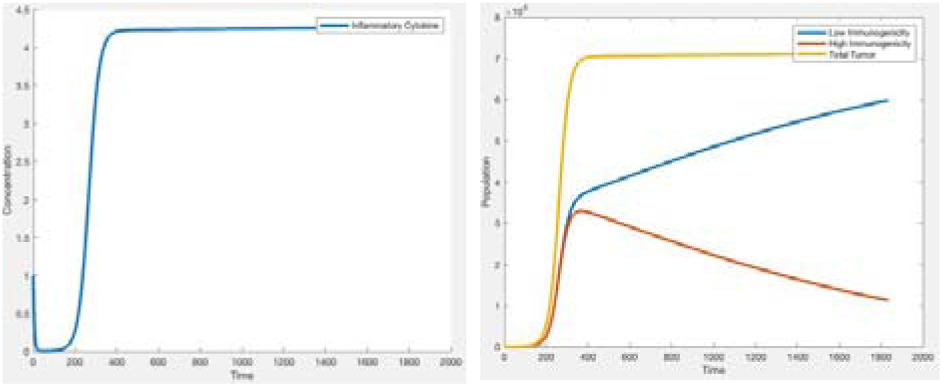
A: Model simulation of inflammatory cytokine concentration without BCG treatment B: Model simulation of low immunogenic cell population, high immunogenic cell population, and total tumor population without BCG treatment Note: The y-axis of each graph is in billions of cells and the x-axis is in days.

In figures 5A and 5B below, the graphs model both the inflammatory cytokines and tumor phenotype populations with BCG treatment. The model was created by adding BCG treatment according to its use in the clinic where it is given every 3-6 weeks. The graphs above show only one BCG treatment cycle and displays the population dynamics associated with it. In Figure 5A, the concentration of inflammatory cytokines increases rapidly until reaching a maximum. The cytokines oscillate once they have reached the highest concentration. In Figure 5B, the low immunogenic population still exceeds the high immunogenic population but at a much more rapid pace. Furthermore, the high immunogenic cells plateau quickly, indicating the strength of the BCG treatment. This causes the bladder cancer tumor population to consist of more low immunogenic cells over time, eventually consisting of almost the entire tumor volume. However, this model may overestimate the effectiveness of the BCG treatment and could be adjusted and calibrated to account for this with actual patient data.

**Figure 5.**
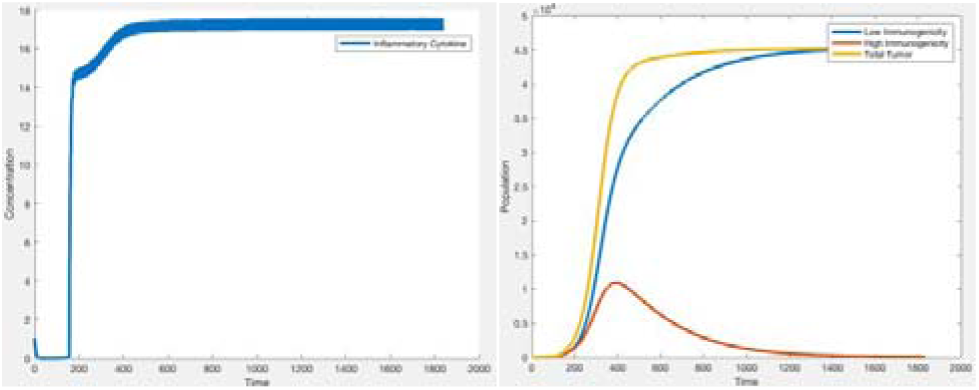
A: Model simulation of inflammatory cytokine concentration with BCG treatment B: Model simulation of low immunogenic cell population, high immunogenic cell population, and total tumor population with BCG treatment

### D. Parameter Scan

After developing the model, parameters were tested to discover which were most sensitive to the entirety of the model with BCG treatment. After testing several different values for each parameter and subsequently graphing the results, it was determined that the most sensitive terms were β, b, and r accordingly. Since β is the scaling term for increasing BCG cytokines, it alters the effectiveness of the BCG treatment, making it most sensitive for tumor cell populations. An example of a parameter scan is shown below, in which β = 10 in Figure 6A and 6B and β = 100 in Figure 6C and D. The graphs reveal that changes in β cause drastic changes to the population dynamics and inflammatory cytokine concentration of the bladder cancer tumor and thus, the entire model.

**Figure 6.**
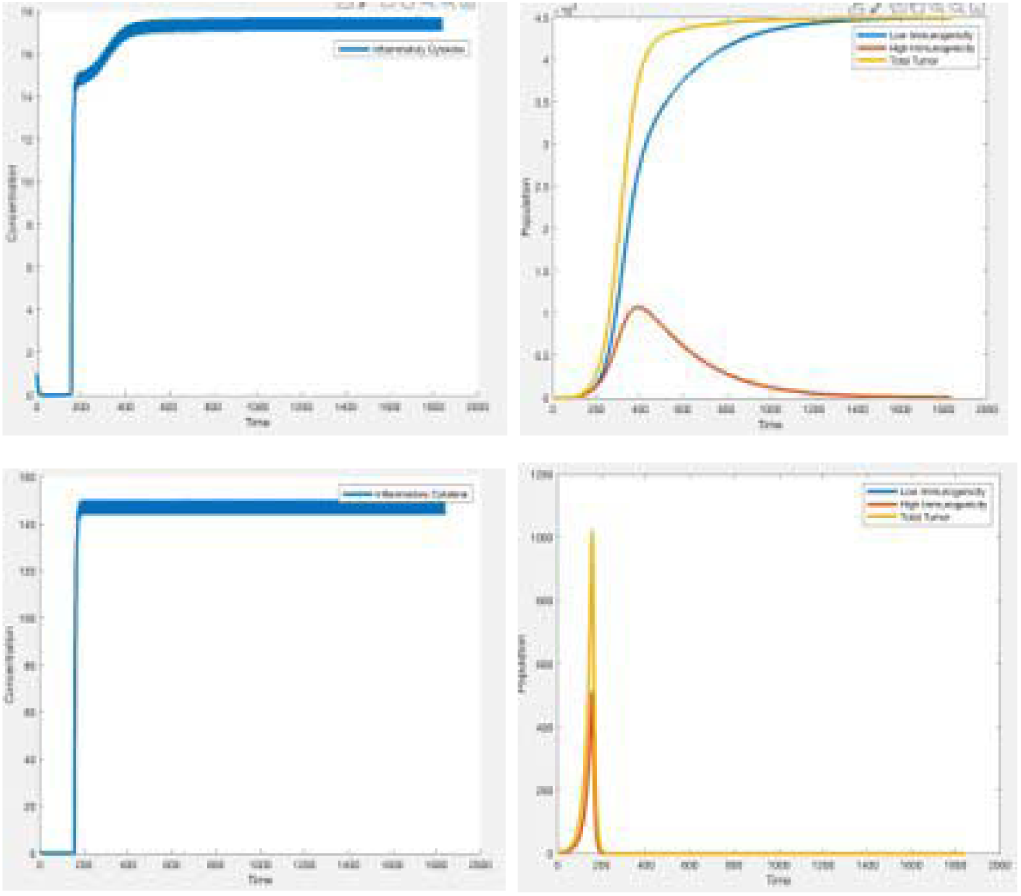
A: Model simulation of inflammatory cytokine concentration with BCG treatment when β = 10 B: Model simulation of low immunogenic cell population, high immunogenic cell population, and total tumor population with BCG treatment when β = 10 C: Model simulation of inflammatory cytokine concentration with BCG treatment when β = 100 D: Model simulation of low immunogenic cell population, high immunogenic cell population, and total tumor population with BCG treatment when β = 100

### E. Varying Initial Conditions

Conventionally, NMIBC cells can be classified into two categories for BCG treatment: BCG refractory or BCG responsive. BCG refractory cells are not ideal and evade treatment, causing the tumor to worsen while BCG responsive cells are ideal for treatment since they respond to the immune system and BCG treatment. The goal for clinicians is to move more bladder cancer cells from BCG refractory to BCG responsive. This can be done by increasing the immunogenicity of these cells through radiotherapy. Therefore, we hypothesized that high immunogenic cells (which are BCG Responsive) make treatment more effective and that radiotherapy could be used to increase immunogenicity and potentially overall tumor response. This relationship is summarized in the graphic (Figure 7) below.

**Figure 7.**
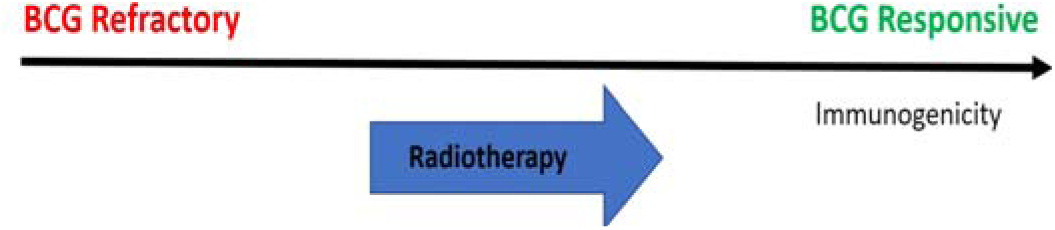
Conventional Treatment Strategies for NMIBC using BCG. By increasing immunogenicity through radiotherapy, more cells become BCG responsive and less cells become BCG refractory.

Thus, this relationship can be affected by the initial phenotypic properties in a tumor. By varying the initial conditions of the model, the impact of immunogenicity and BCG responsive and refractory cells can be determined. The graphs below indicate what occurs when the initial number of low immunogenic cells is less than, equal to, and greater than the number of high immunogenic cells. As the graphs reveal, initial conditions can greatly affect tumor phenotype populations by making them grow at faster or slower rates.

**Figure 8:**
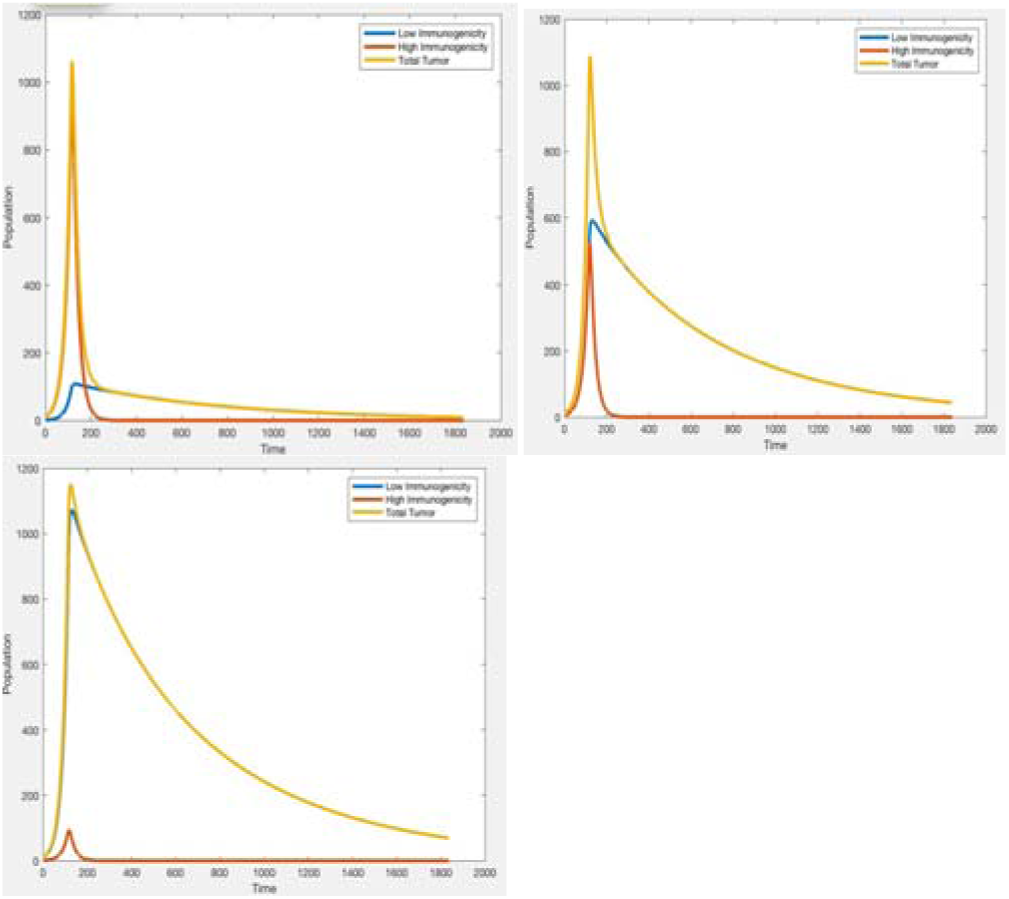
A: Model simulation of low immunogenic cell population, high immunogenic cell population, and total tumor population when the number of low immunogenic cells is less than the number of high immunogenic cells B: Model simulation of low immunogenic cell population, high immunogenic cell population, and total tumor population when the number of low immunogenic cells is equal to the number of high immunogenic cells C: Model simulation of low immunogenic cell population, high immunogenic cell population, and total tumor population when the number of low immunogenic cells is greater than the number of high immunogenic cells

### F. Stability Analysis

In order to determine the parameters needed to create a stable model, the differential equations were placed into Mathematica and solved. Furthermore, the Eigenvalues were computed for each equation to give a set of rules for the parameters to follow. Thus, in order for the model to be stable with the high immunogenic cells outcompeting, the following three conditions must be met:

1. r > d The growth rate must exceed the death rate.
2. b > b_2_ Death due to cytokines of low immunogenic cells exceeds that of high immunogenic cells
3. > 4(r-d) Cytokine decay rate exceeds four times the growth rate minus death rate or natural growth rate

## IV. Discussion and future work

Based on the results of the balder cancer ODE models, it can be concluded that phenotypes and immunogenicity affect tumor growth and treatments. Furthermore, it may be possible to recommend certain treatments based on the phenotypic properties of the tumor cells. This would be possible if patient data was incorporated into the model in order to predict outcome based on the initial phenotypes. In the future, we would like to expand this model in order to include not only immunogenicity but also proliferation to accurate predict patient response to BCG treatment in NMIBC.

## Acknowledgements

I would like to thank the entire Integrated Mathematical Oncology department at H. Lee Moffitt Cancer Center and Research Institute and my mentors for making this research possible as a high school student. Furthermore, this research is the property of H. Lee Moffitt Cancer Center and Research Institute. I would especially like to thank my mentors during this program who played a key role in helping me perform this research: Dr. Meghan Ferrall-Fairbanks, Dr. Philipp Altrock, and Dr. Gregory Kimmel.

